# All-Resolutions Inference for Brain Imaging

**DOI:** 10.1101/226126

**Authors:** Jonathan D. Rosenblatt, Livio Finos, Wouter D. Weeda, Aldo Solari, Jelle J. Goeman

**Affiliations:** Department of IE&M and Zlotowsky Center for Neuroscience, Ben Gurion University of the Negev, Israel; DPSS, University of Padua, Italy; Methodology & Statistics Unit, Department of Psychology, Leiden University, Leiden, The Netherlands, and Leiden Institute for Brain and Cognition, Leiden University, Leiden, The Netherlands; DEMS, University of Milano-Bicocca, Italy, and NeuroMI - Milan Center for Neuroscience, University of Milano-Bicocca, Italy; Department of Medical Statistics and Bioinformatics, Leiden University Medical Center, The Netherlands

## Abstract

The most prevalent approach to activation localization in neuroimaging is to identify brain regions as contiguous supra-threshold clusters, check their significance using random field theory, and correct for the multiple clusters being tested. Besides recent criticism on the validity of the random field assumption, a spatial specificity paradox remains: the larger the detected cluster, the less we know about the location of activation within that cluster. This is because cluster inference implies “there exists at least one voxel with an evoked response in the cluster”, and not that “all the voxels in the cluster have an evoked response”. Inference on voxels within selected clusters is considered bad practice, due to the voxel-wise false positive rate inflation associated with this circular inference. Here, we propose a remedy to the spatial specificity paradox. By applying recent results from the multiple testing statistical literature, we are able to quantify the proportion of truly active voxels within selected clusters, an approach we call All-Resolutions Inference (ARI). If this proportion is high, the paradox vanishes. If it is low, we can further “drill down” from the cluster level to sub-regions, and even to individual voxels, in order to pinpoint the origin of the activation. In fact, ARI allows inference on the proportion of activation in all voxel sets, no matter how large or small, however these have been selected, all from the same data. We use two fMRI datasets to demonstrate the non-triviality of the spatial specificity paradox, and its resolution using ARI. One of these datasets is large enough for us to split it and validate the ARI estimates. The conservatism of ARI inference permits circularity without losing error guarantees, while still returning informative estimates.

## 2 Introduction

The fundamental building block of brain mapping with functional magnetic resonance imaging (fMRI) is arguably the localization of evoked brain responses to cognitive stimuli. Localization is typically performed by correlating a sequence of stimuli to the sequence of measured blood oxygenation levels (BOLD) at each brain region, and then testing for the statistical significance of these correlations. Correlation should be understood in a broad sense, and may involve simple correlations, linear models, non-linear models, machine learning classifiers, and more. A region is declared “active”, or “information-encoding”, if this correlation is statistically significant compared to an “inactive region” null hypothesis. Clearly, testing many regions in the brain introduces a severe multiplicity problem, leading, for example, to the detection of information-encoding regions in dead salmon fish [Bennett et al., 2009]. Error rate inflation was acknowledged by the neuroimaging community early on, and led to an awareness in the community of the dangers of *selective inference*. Selective inference includes *selective testing*, better known as *multiple testing* [Friston et al., 1991, Genovese et al., 2002], and also *selective estimation*, better known as *voodoo correlations, circular inference*, and *double-dipping* [Vul et al., 2009, Kriegeskorte et al., 2009, Rosenblatt and Benjamini, 2014]. The community’s awareness of selective inference is manifested in the fact that all software suites for brain imaging (SPM, FSL, BrainVoyager, and AFNI) include several multiplicity correction methods. It is also manifested in the fact that it is impossible to publish a paper in the field if multiplicity has not been addressed.

The localization of activation in the brain requires the neuroscientist to choose the type of inference to make. This includes (i) the scale of brain regions, and (ii) the choice of error guarantees. The scale of brain regions may vary from a single volume element (voxel), to multiple contiguous elements defined by their shape, their anatomical properties, or their functional properties. These are known as *voxel-wise* inference, *searchlight, anatomical regions of interest* (ROIs), and *functional regions of interest*, respectively. The error guarantee applied is typically the *family-wise error rate* (FWER), or the *false discovery rate* (FDR). FWER is interpretable as the proportion of studies in which false discoveries are made, and FDR as the average *false discovery proportion* over all studies. Historically, the first inferences were voxel-wise, or anatomical ROIs, with FWER error guarantees. Then came functional ROIs, FDR controls, multivariate searchlights, and others. Today, combinations of all scales of inference with all error controls can be found in the neuroimaging literature [e.g. Poldrack et al., 2011].

A scale of inference which deserves particular attention is *cluster-based inference*. The idea of cluster inference dates back to Poline and Mazoyer [1993], Forman et al. [1995], and Friston et al. [1996]. It is now the most common type of inference, being the default option in several popular software suites. Cluster inference can be seen as inference at a *data-driven scale*. This is because the size of the clusters is not selected a priori, but rather determined by the data used for inference. The fact that clusters are both defined and tested with the same data introduces a statistical circularity challenge typically solved using a *random field theory* (RFT) approach, which permits both FWER control on clusters [Taylor and Worsley, 2007a], and FDR control on clusters [Chumbley et al., 2010].

Unfortunately, cluster inference has been heavily criticized, firstly for inappropriate error guarantees, as recently shown in the high-profile contribution of Eklund et al. [2016].

Cluster inference also suffers from low spatial resolution [Woo et al., 2014], which is demonstrated by the following paradox. Since discovering a cluster means that “there exists at least one voxel with an evoked response in the cluster”, and not that “all the voxels in the cluster have an evoked response”, it follows that *the larger the detected cluster, the less information we have on the location of the activation*. Moreover, cluster-based inference gives no information on the extent of the activation within the cluster.

The matter of low spatial resolution can be remedied by a hierarchical approach^1^—a “drill-down” from discovered clusters to subsets of those clusters, and ultimately, to the voxel level. The aim of a drill-down is to localize the activation within the cluster, and to quantify its extent. However, it is typically a forbidden practice, because voxel-wise error guarantees will not hold when inferring on voxels within selected clusters. Such a drill-down would entail three layers of circularity: creating clusters, inferring on clusters, and inferring on voxels within clusters. Acknowledging the three layers of circularity for valid inference is a formidable mathematical challenge. The purpose of this manuscript is to report the application of a recent advance in the hierarchical inference literature, namely that of Goeman and Solari [2011], to permit valid circular inference of this type in neuroimaging.

The *All-Resolutions Inference* (ARI) of Goeman and Solari [2011] allows more than a single drill-down from the cluster to the voxel: it allows the researcher to apply any data-driven region selection and estimate the *proportion of true discoveries* (PTD^2^) of any subregion—clusters in our case—all from the same dataset. ARI accounts for the circularity by controlling the FWER over all possible subsets of the brain, large or small, contiguous or non-contiguous, using *closed testing* [Marcus et al., 1976]. Extreme conservatism for FWER control over these exponentially many regions can be avoided, if tests for overlapping regions are highly correlated. ARI exploits the powerful Simes test, which exhibits the necessary correlation structure. The Simes test is valid under the assumption of the Simes inequality, which is implied by the *positive regression dependency on subsets* condition (PRDS), established for brain maps by Nichols and Hayasaka [2003]. The assumption of the Simes inequality is well-known, since it is also necessary for the FDR-controlling procedure of Benjamini and Hochberg [1995]. The closed testing procedure, in combination with the Simes inequality, guarantees FWER control on all statements made using ARI, which means that with probability at least 95% no region has an overestimated PTD.

In Section 3, we prove that ARI returns lower bounds on the PTD, at all scales simultaneously, and for various selection criteria. Examples include voxels within clusters, voxels within searchlights, anatomical ROIs within functional ROIs, etc. The non-technical reader may want to skip directly to Section 4, where we apply ARI to several datasets. From these we learn that the spatial specificity paradox exists empirically and cannot be ignored. For some datasets and thresholds, clusters consist of mostly active voxels, while for other datasets and thresholds, clusters consist of mostly inactive voxels. For the latter clusters, ARI allows one to look at data-driven subclusters to better pinpoint the location of active voxels.

While not our initial motivation, ARI may also serve for cluster inference, thus replacing the RFT *p*-values. Since ARI does not rely on RFT, it eschews the inaccuracies of cluster inference recently reported by Eklund et al. [2016], and it avoids the computational burden of resampling-based inference. We elaborate on cluster selection with ARI in Section 5.1.

## 3 The All-Resolutions Inference Framework

We start with an exposition of the ARI method, the datasets to which it has been applied, and the manner in which it has been applied.

### 3.1 Overview of the Framework

The brain *B* is a collection of *m* voxels. We assume that a test statistic for activation has been calculated for each voxel, which can be converted into a voxel-wise *p*-value.

Researchers are interested in inference on subsets of the brain. In general, we use the term *voxel set* for any subset of the brain, possibly non-contiguous. Special types of voxel sets are regions, clusters, and searchlights. We denote 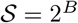 as the collection of all 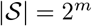 voxel sets, where |·| denotes the cardinality of a set. Brain regions are interesting if they contain many truly active voxels. Let the unknown voxel set *A* ⊆ *B* be the set of all *truly* active voxels. For any voxel set 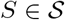, denote the number of truly active voxels *a*(*S*) = |*A* ∩ *S*|, and their proportion (PTD) by *π*(*S*) = *a*(*S*)/|*S*|.

ARI uses the methods of Goeman and Solari [2011] and Meijer et al. [2016] to construct lower confidence bounds 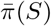 for the set-wise proportion of active voxels, simultaneously for all possible sets. The (1 – *α*) lower confidence bound is such that

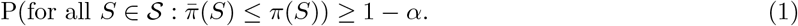

Simultaneity over all 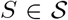, i.e. the fact that the “for all” statement is inside the probability statement, crucially makes all inference based on 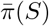 robust against circular selection of sets. With probability at least 1 – *α* the bound is valid for all *S*, and therefore for one or more selected *S*, regardless of how they were selected. Simultaneity, in turn, implies FWER control over all statements made about the selected *S*.

In particular 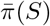 may also be calculated for sets of one voxel, for which it takes the values 0 or 1. In ARI the singleton sets for which 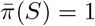 precisely correspond to the voxels rejected by the procedure of Hommel [1988], a uniform improvement of Bonferroni. As we shall see below, however, ARI is more powerful for larger sets than for small ones, and may give large values of 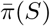 even if no voxel in *S* is significant by Hommel [1988].

### 3.2 Simes Test and Simes Inequality

To derive (1) we start by defining for every voxel set 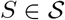 the null hypothesis

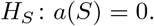

*H_S_* is the usual null hypothesis for cluster-wise inference: rejecting *H_S_* indicates that there is at least one active voxel in *S*. We test every *H_S_* with the Simes test [Simes, 1986], rejecting *H_S_* at level *α* if and only if *p_S_* ≤ *α*, where

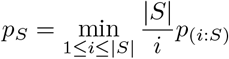

and *p*_(*i:S*)_ is the *i*th smallest *p*-value among voxels in *S*.

The Simes test is valid if P(*p_S_* ≤ *α*) ≤ *α* for all *S* for which *H_S_* is true. For the validity of the ARI procedure as a whole, however, we only need this to hold for the set *S* = *B* \ *A* of all non-active voxels, the largest set for which *H_S_* is true. We assume that

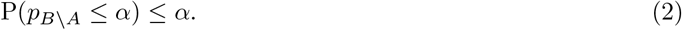

Equation (2), the Simes inequality, is the most important assumption required for ARI. The assumption of the Simes inequality is frequently made in the multiple testing literature, and oft-used procedures such as those of Hommel [1988], Hochberg [1988] and Benjamini and Hochberg [1995] make the same assumption. There is, therefore, much ongoing research on sufficient conditions for the validity of the Simes inequality [Finner et al., 2014]. It has been shown to hold for independent *p*-values, and under various conditions implying non-negative correlations between *p*-values, one of which is the PRDS condition. Nichols and Hayasaka [2003] have shown that PRDS, and therefore the Simes inequality, is valid for brain maps.

### 3.3 All-Region FWER Control

The tests for the 2^*m*^ hypotheses *H_S_*, 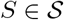, must be corrected for multiple testing. A powerful method for this is closed testing [Marcus et al., 1976]. Intuitively, closed testing means that if a particular configuration of true and false null hypotheses may inflate false-positive rates, then the rates for this configuration should be controlled explicitly, for all possible configurations. Formally, in closed testing a hypothesis *H_S_* is rejected if and only if *H_I_* is rejected for all *I* ⊇ *S*. Closed testing controls the FWER at level *α* for all *H_S_*, 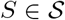, under the simple condition that *H_B\A_* is a valid *α*-level test, i.e. under the assumption of the Simes inequality.

Meijer et al. [2016] have proven that closed testing with Simes tests rejects a hypothesis *H_S_* if and only if

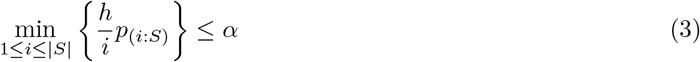

where

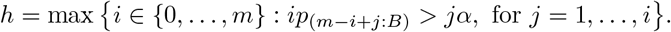

Note that *h* depends on *α* and on the *p*-values of voxels outside *S*. Closed testing is a very powerful procedure, and the cost of FWER control over all possible hypotheses is therefore relatively light, taking into account the fact that 2^*m*^ hypotheses are tested; compared to the unadjusted Simes test, the critical value *α* is multiplied only by a factor *h*/|*S*| ≤ *m*/|*S*|. This is because the Simes inequality condition ensures that even though many comparisons are considered, test statistics of overlapping regions are highly correlated and the distribution of the maximal *pJ* over the supersets *J* is “tight”. The calculation of FWER-adjusted *p*-values, for any *S*, is described in Meijer et al. [2016].

### 3.4 Proportion of Truly Active Voxels (PTD)

Lower confidence bounds for the percentage of truly active voxels (PTD above) follow from the result of the closed testing procedure by the argument given by Goeman and Solari [2011]: specifically, if for some *k* ≥ 0, *H_I_* is false for all subsets *I* ⊆ *S* with |*I*| = |*S*| – *k*, then there is at least one active voxel in each such *I*, and therefore there are at least *k* + 1 active voxels in *S*. Goeman and Solari [2011] defined 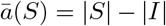, where *I* is the largest subset of *S* such that *H_I_* was not rejected by the closed testing procedure.

That 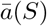 is a simultaneous lower bound on the PTD of the region, i.e. that

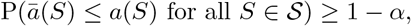

follows immediately from the FWER-control of the closed testing procedure, and (1) follows immediately by setting 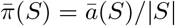.

For the case of Simes tests, we have

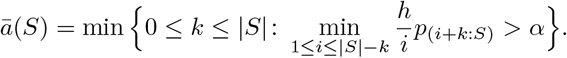

Efficient ways to calculate this quantity are given by Meijer et al. [2016]. The lower bound is the minimum number of *p*-values that can be removed so that (3) is violated for the resulting subset.

We note that by the properties of closed testing 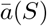 never increases when drilling down, i.e. reducing *S* to a subset. 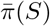, however, may increase when drilling down unless 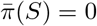. It may pay to drill down for PTD, but never for regions where no signal is found.

### 3.5 Image Analysis Pipeline

ARI brings unprecedented freedom in looking at the data, choosing regions in any desired way, calculating PTDs for the chosen regions, and possibly reconsidering the selection, e.g. if regions are too small or too large, or have a disappointingly low percentages of active voxels. Different criteria may be used to select different regions. FWER control is guaranteed as long as the *α*-level and the method of calculating *p*-values have been decided before seeing the data.

To demonstrate the spatial specificity paradox, and its resolution with ARI, we start by selecting clusters with a standard analysis pipeline, and then compute PTDs in these clusters. Given a Z-score map, we defined clusters of interest using pre-specified cluster-forming *Z*-threshold and minimal cluster size. The cluster size threshold we use is equivalent to an RFT significance threshold. The latter choice is only for conforming to current practice. We emphasize that ARI estimates do not rely, in any way, on RFT inference. RFT significance and ARI significance do not necessarily coincide: there may be regions significant under RFT but not under ARI, and vice versa.

To pinpoint activation within these clusters, we drilled down to smaller regions by increasing the cluster-forming threshold to *Z* > 4 and looking at the clusters, significant by RFT or not, that are contained within the significant clusters at *Z* > 3.2.

The cluster-forming thresholds of 3.2 and 4 are arbitrary, and we did not fix them before seeing the data. In ARI this post hoc choice of regions, and the manner in which they are selected, does not invalidate FWER control for inference on the chosen regions.

For convenience, we collect our two selection criteria as used in this manuscript:

1. *Z* > 3.2 clusters with a size/significance threshold.
2. *Z* > 4 clusters which lie within *Z* > 3.2 clusters.

### 3.6 fMRI Data

To demonstrate the results of ARI, we applied it on two fMRI datasets we term *Go*/*No-go*, and *Auditory*. The Go/No-go dataset consists of 34 subjects performing an emotional go/no-go task [Lee et al., under review]. Participants had to press a button when presented with faces with a certain emotional expression (go condition), and withhold their response to faces with a neutral expression (no-go condition). The go and no-go conditions were then reversed, to avoid confounding with button-press-related activation. The Auditory dataset was collected by Pernet et al. [2015], and generously shared via the OpenfMRI initiative at https://openfmri.org/. It consists of 218 subjects passively listening to vocal (i.e. speech) and non-vocal sounds. The large dataset allowed us to validate our PTD estimates on different subjects. We used two mutually exclusive sets, an original sample with a typical fMRI sample size of 33 subjects, and a validation sample of 66 subjects, serving as a “ground-truth”. We used only 66 and not the 185 = 218 – 33 remaining subjects, so that the “ground-truth” is not driven by infinitesimally small effects.

## 4 Results

### 4.1 Go/No-go Data

Group analysis of the No-go > Go contrast highlighted 11 regions of interest commonly found in studies using the Go/No-go paradigm (see Figure 1).

**Figure 1:**
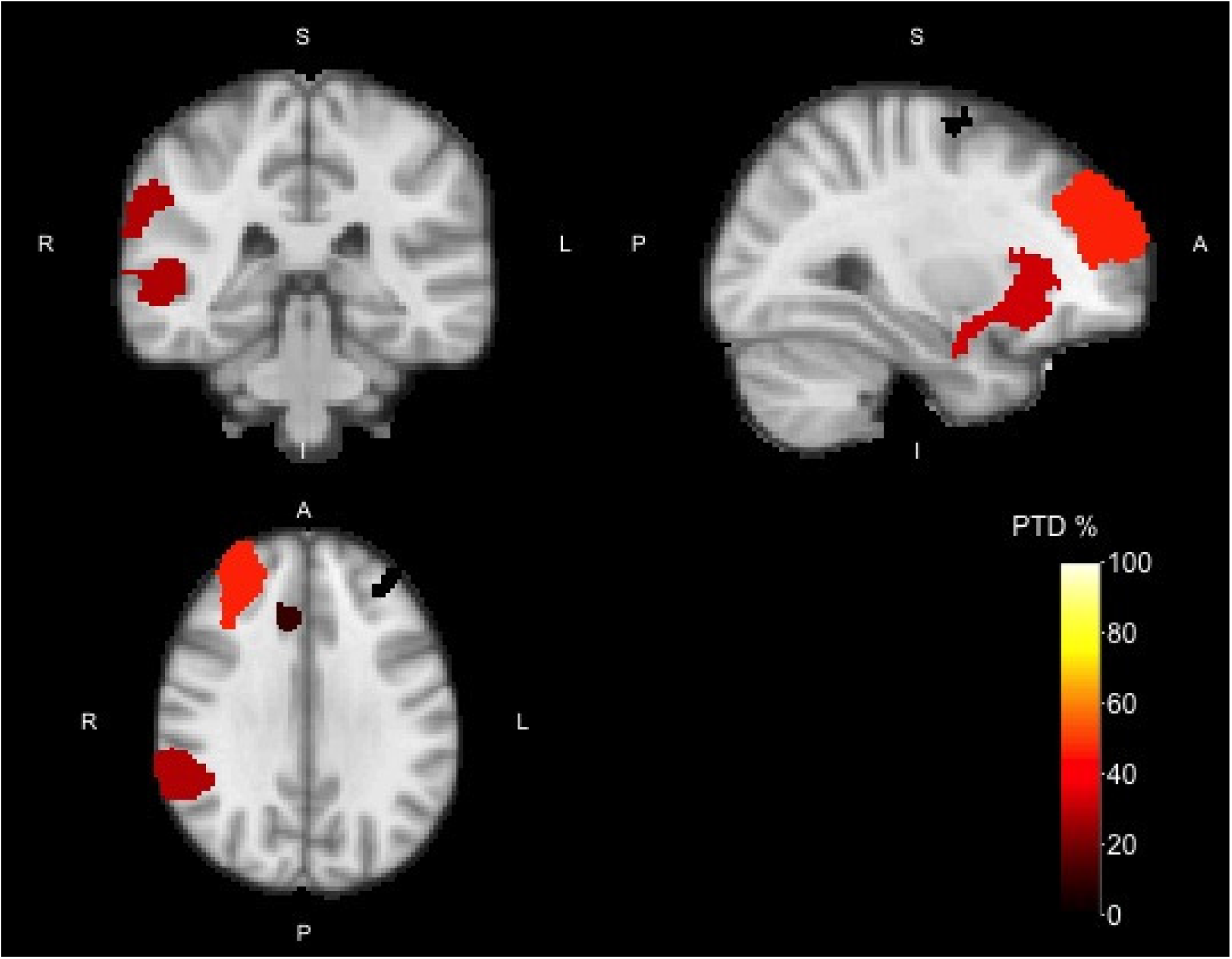
Activation map of the No-go > Go contrast with cluster-forming threshold *Z* > 3.2 for the Go/No-go data. Colors indicate the PTD for each of the clusters.

These regions included the left and right insular cortex (IC) extending into the frontal orbital cortex (FOC), the left and right frontal pole (FP), the right middle (MTG) and superior temporal gyrus (STG) extending into the angular gyrus (AG), the right (para)cingular gyrus (PCG), right superior frontal gyrus (SFG), right precuneus, and the right precentral gyrus.

PTDs for these ROIs were highest for the region spanning the right MTG, STG and angular gyrus (2191 voxels, PTD = 28.5%), the region spanning the right frontal pole (1845 voxels, PTD = 46.2%), and the region spanning the right insular cortex and frontal orbital cortex (1400 voxels, PTD = 32.4%). The regions spanning the left insular cortex and frontal orbital cortex (421 voxels, PTD = 5.9%), and (para)cingular gyrus (304 voxels, PTD = 10.9%) had lower PTDs, while for the other regions (right SFG, precuneus, precentral gyrus, and left frontal pole) the PTD was 0%.

Of the regions with a PTD of 0%, which are not expected to have many active voxels, all four regions had significant RFT-corrected *p*-values: The right SFG (698 voxels, RFT *p* < .001, ARI *p* = .068, PTD = 0%), right precuneus (245 voxels, RFT *p* = .010, ARI *p* = .533, PTD = 0 %), right precentral gyrus (232 voxels, RFT *p* = .012, ARI *p* = .069, PTD = 0%), and left FP (698 voxels, RFT *p* = .029, ARI *p* = .497, PTD = 0%).

Details of the clusters can be found in Table 1. Details include the name and size of the clusters, ARI-estimated number and proportion of active voxels, and ARI *p*-value. In addition we include *standard* details on the location of the cluster maximum (MNI coordinates), *Z*-value of the maximum and RFT-corrected *p*-values for each cluster.

**Table 1:**
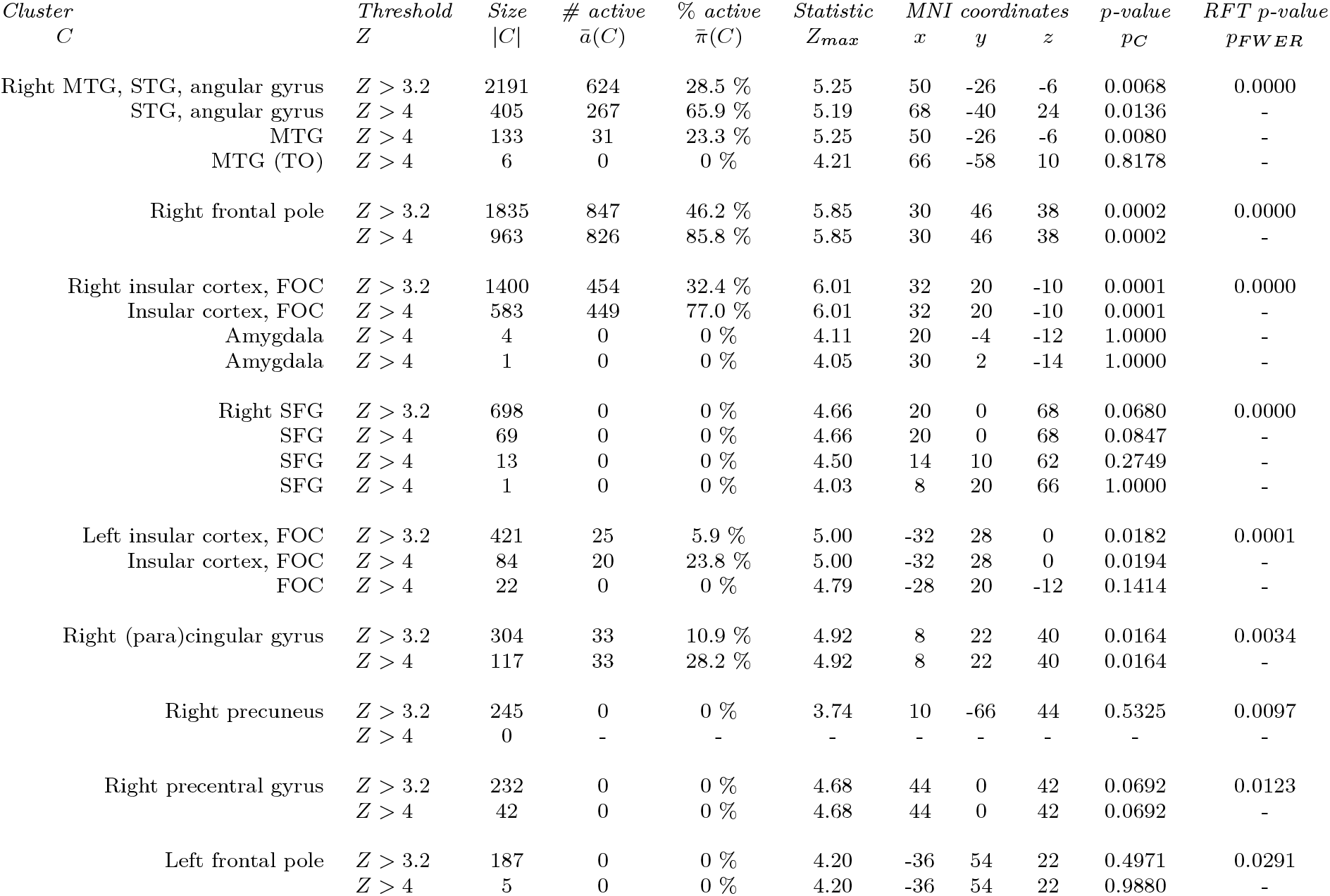
Go/No-go data: clusters identified with threshold *Z* > 3.2 (RFT *p* < .05, cluster size = 161), with “drill down” clusters at *Z* > 4.

To pinpoint the location of the truly active voxels, i.e. to “drill down”, we could infer on all the voxels in selected clusters. ARI guarantees that this inference would be valid but, alas, low-powered. Alternatively, we may increase the cluster-forming threshold. Inference would still be valid, and the PTD would increase, until all supra-threshold voxels are truly active.

Using a cluster-forming threshold of *Z* > 4 and looking at these “drill down” clusters within the *Z* > 3.2 clusters, we obtained 19 clusters (Figure 2 and Table 1). The cluster spanning the right MTG, STG, and AG (PTD = 28.5%), is now divided into three smaller subclusters: STG/AG (405 voxels, PTD = 65.9%), MTG (133 voxels, PTD = 23.3%) and a small cluster also in the MTG (6 voxels, PTD 0%). The division in multiple subclusters and their corresponding PTD is consistent with areas often indicated in inhibition studies (Neurosynth meta-analysis [Yarkoni et al., 2011], keyword ‘nogo’). The subclusters with high a PTD (STG/AG) are often found in inhibition studies, while the chance of finding a cluster in the MTG is much smaller.

**Figure 2:**
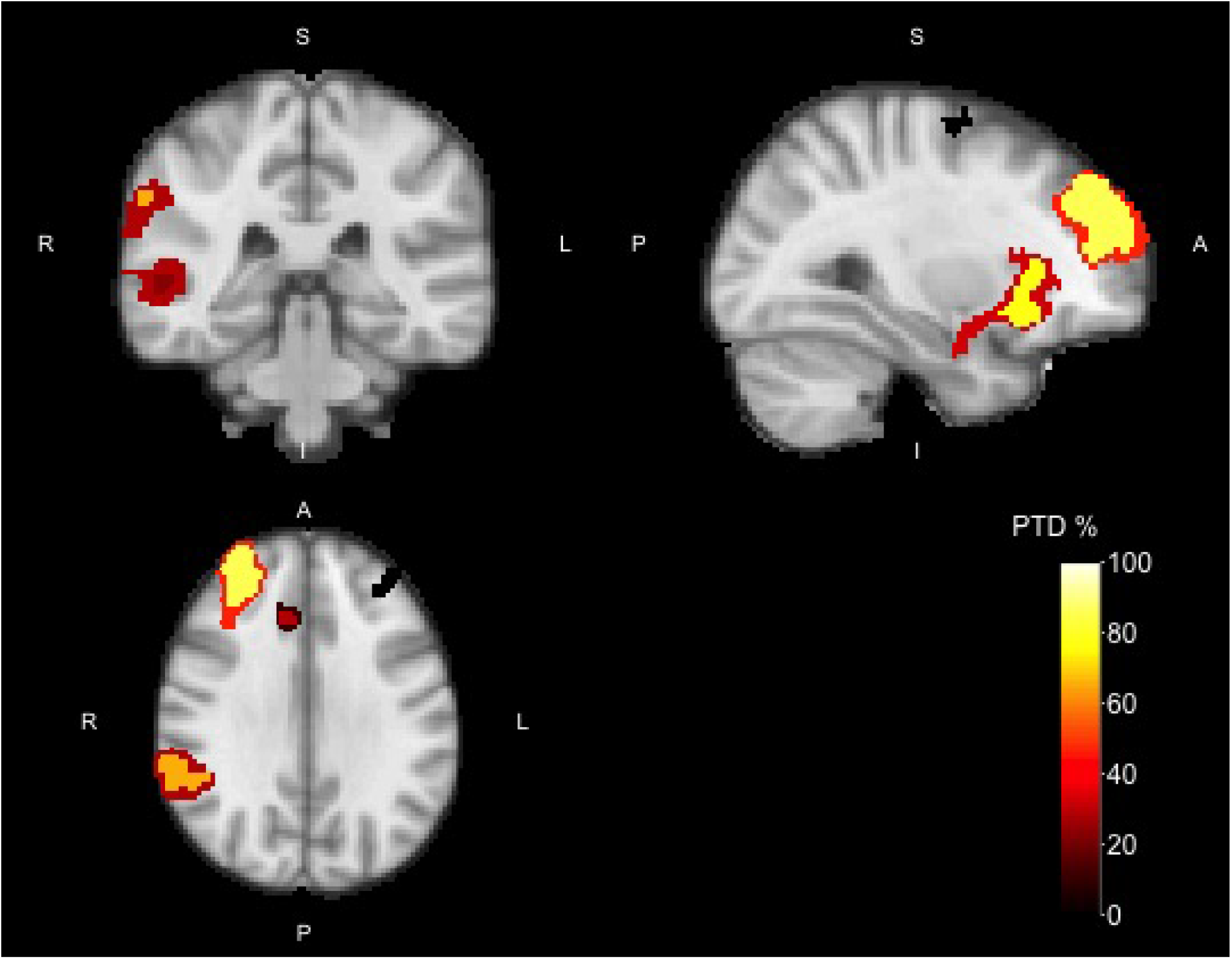
Activation map of the No-go > Go contrast with cluster-forming threshold *Z* > 4 for the Go/No-go data overlaid to the *Z* > 3.2 map. Colors indicate the PTD for each of the clusters.

The right FP contained only one smaller cluster (963 voxels, PTD = 85.8%) with a high PTD. The region spanning the right IC and FOC also was divided into three subclusters: one spanning the IC and FOC (583 voxels, PTD = 77.0%) and two spanning the amygdala (1 and 4 voxels respectively, PTD = 0%). The left IC/FOC cluster contained two small clusters, one spanning the IC/FOC (84 voxels, PTD = 23.8%) and one spanning only the FOC (22 voxels, PTD = 0%). Again, these results are consistent with the literature regarding inhibition studies, where the IC/FOC cluster is more often found than the FOC cluster (Neurosynth meta-analysis [Yarkoni et al., 2011], keyword ‘nogo’).

The right PCG contained one small cluster (117 voxels, PTD = 28.2%). For the right SFG, drilling down revealed three clusters all within the SFG (69, 13, and 1 voxel), all with a PTD of 0%. The other regions contained no active voxels in the smaller regions.

The ARI drill-down analysis is thus consistent, and more informative about the smaller clusters. As evident from the drill-down analysis, the new smaller region in the right FP has a higher PTD (85.8% vs 46.2%), indicating that this smaller area contains most relevant information. The same holds for the right IC/FOC (32.4% vs 77.0%) and STG/AG areas (28.2% vs 65.9%), where the new smaller regions now contain a more acceptable number of truly active voxels, and the spatial specificity paradox is alleviated. Above all, drilling down to smaller clusters reveals the clusters that are interesting (i.e. contain active voxels), and those which can be discarded.

### 4.2 Auditory Data

#### 4.2.1 Inference

Group analysis on the first set of 33 subjects of the Vocal > Non-vocal contrast showed activity in 6 regions of interest commonly found in auditory studies. Details of the clusters can be found in Table 2. We observed activity bilaterally in the superior temporal gyrus (STG), planum temporale (PT), Heschl’s gyrus (HG), inferior frontal gyrus (IFG), and amgydala, and activity in the right precentral gyrus.

**Table 2:**
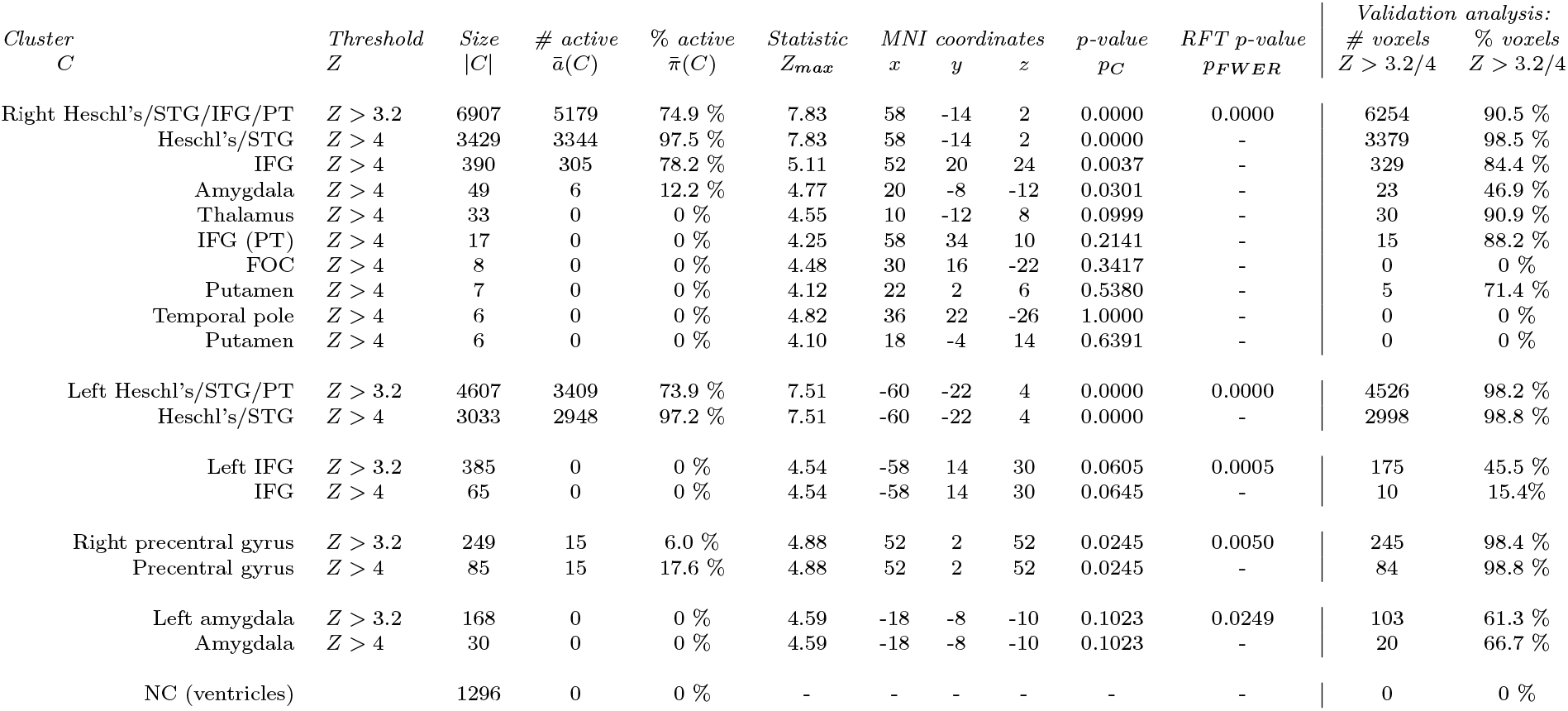
Auditory data: clusters identified with threshold *Z* > 3.2 (RFT *p* < .05, cluster size = 118), with “drill down” clusters at *Z* > 4.

As can be seen in Figure 3, the activity in the right hemisphere covered one large cluster (6907 voxels), with a PTD of 74.9% (with exception of the precentral gyrus; 249 voxels, PTD = 6.0%). In the left hemisphere these same areas were divided amongst three regions: HG/STG/PT (4607 voxels, PTD = 73.9%), IFG (385 voxels, PTD = 0%), and the amygdala (168 voxels, PTD = 0%).

**Figure 3:**
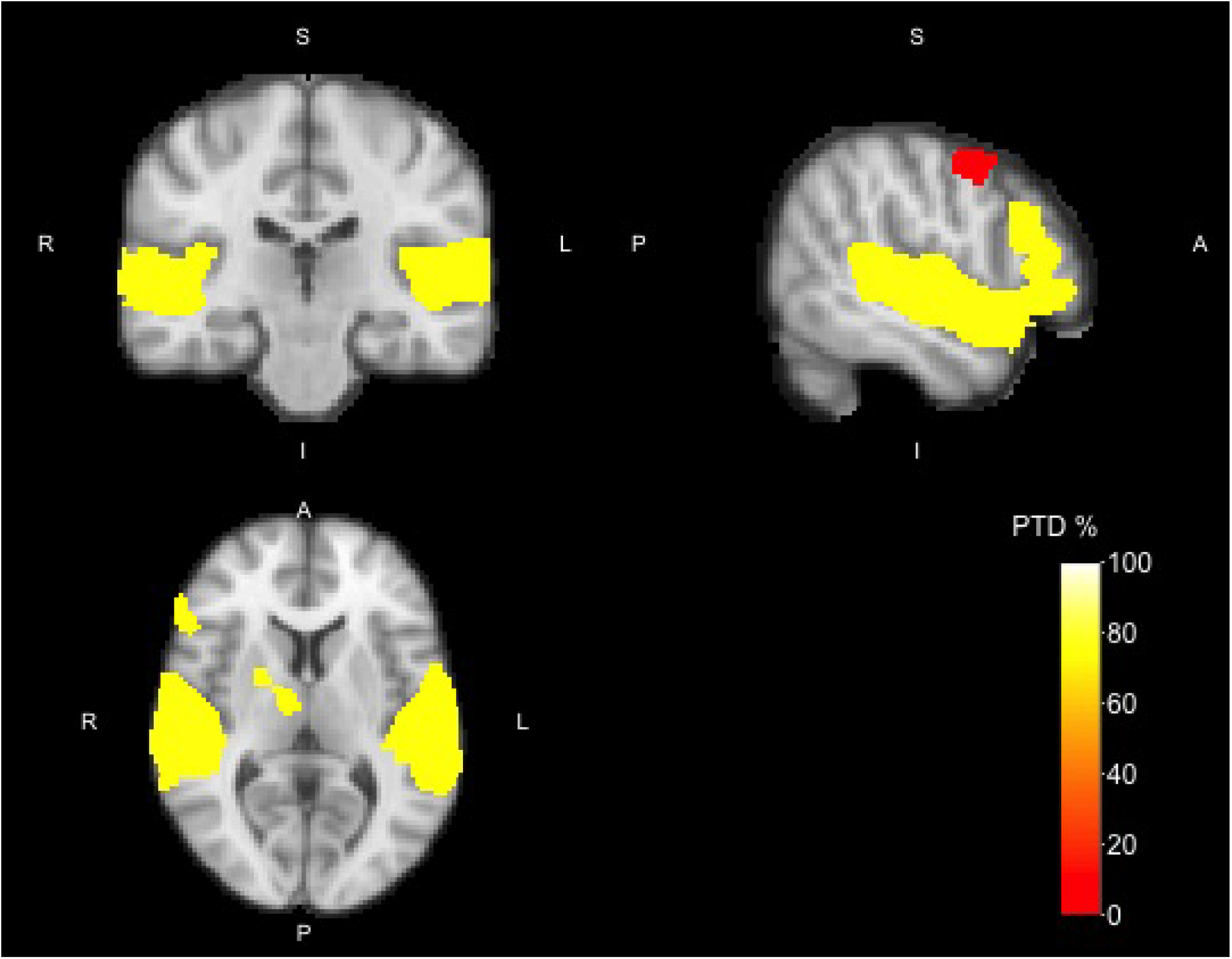
Activation map of the Vocal > Non-vocal contrast with cluster-forming threshold *Z* > 3.2 for the Auditory data. Colors indicate the PTD for each of the clusters.

As with the previous analysis we can now drill down with a higher cluster-forming threshold to check the proportion of active voxels in the smaller regions. With a cluster-forming threshold of *Z* > 4, we now see 15 smaller regions within the *Z* > 3.2 clusters (see Figure 4). The large cluster in the right hemisphere of the temporal cortex in the *Z* > 3.2 analysis now separates into 9 separate clusters. Three of these clusters had a PTD larger than 0%: HG/STG (3429 voxels, PTD =97.5%), IFG (390 voxels, PTD = 78.2%), and the amygdala (49 voxels, PTD = 12.2%); the other clusters contained no active voxels (see Table 2 for details). The HG/STG cluster in the left hemisphere was now also smaller, with a high PTD (3033 voxels, PTD = 97.2%). The smaller clusters in the left amygdala, and IFG, contained no active voxels (PTD = 0%). The smaller cluster in the right precentral gyrus (85 voxels, PTD = 17.6%), had a slightly higher PTD value.

**Figure 4:**
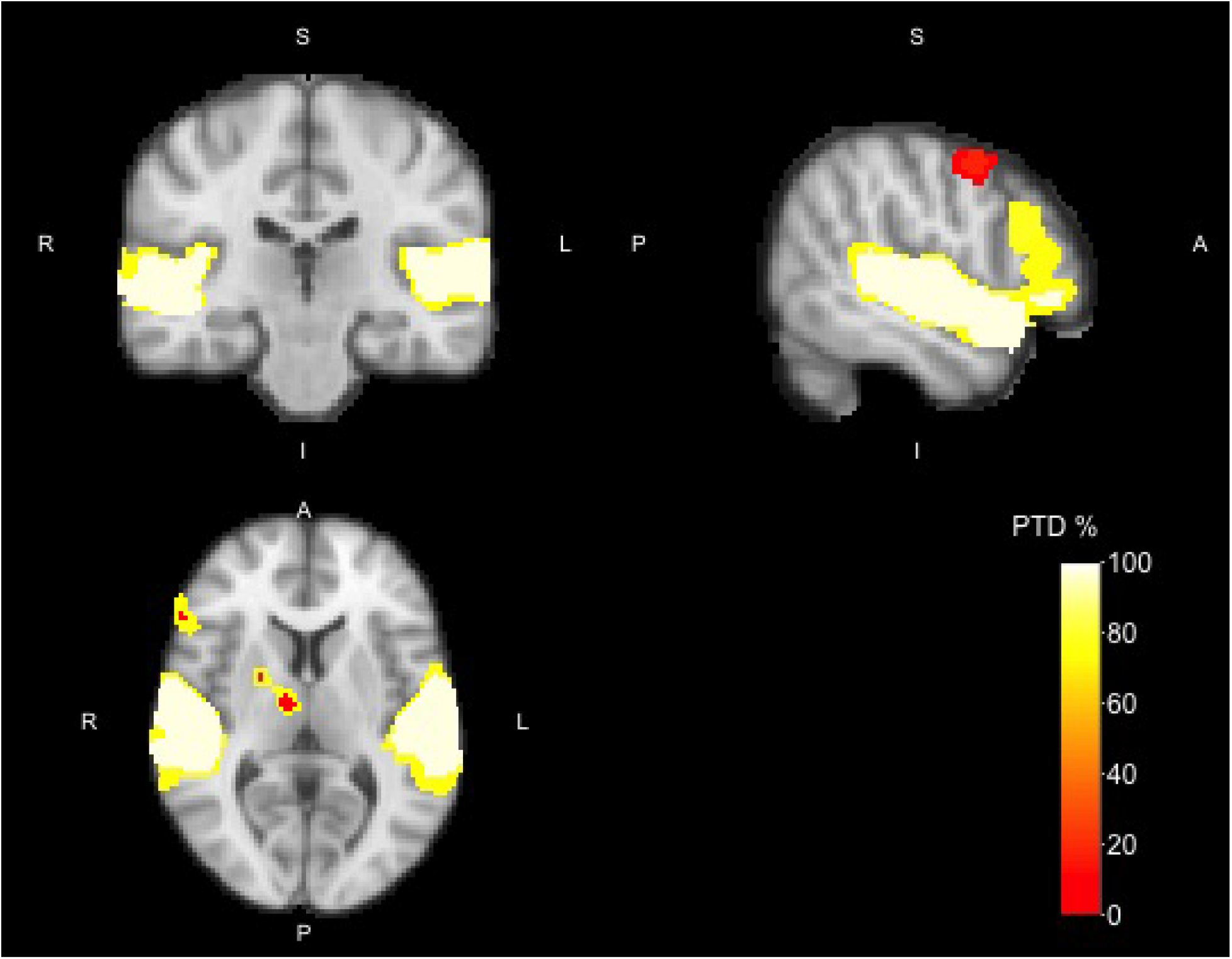
Activation map of the Vocal > Non-vocal contrast with cluster-forming threshold *Z* > 4 for the Auditory data overlaid to the *Z* > 3.2 map. Colors indicate the PTD for each of the clusters.

#### 4.2.2 Validation

To check the validity of the ARI method we used the *Z* > 3.2 and *Z* > 4 clusters from the previous analysis and calculated the number of supra-threshold voxels of these clusters using data from the second set of 66 subjects. The following results confirms that the ARI PTD bounds are both informative and statistically valid, despite the circular analysis.

For *Z* > 3.2, the validation dataset showed 90.5% supra-threshold voxels in the right Heschl’s gyrus/STG, compared to a PTD of 74.9% in the original sample. In the left Heschl’s gyrus/STG we found 98.2% supra-threshold voxels, compared to a PTD of 73.9% in the original sample. In the left IFG the number of supra-threshold voxels was 45.5%, compared to a PTD of 0% in the original sample. The right precentral gyrus showed 98.4% supra-threshold voxels, compared to a PTD of 6.0% in the original sample. In the left amygdala we observed 61.3% supra-threshold voxels, with a PTD of 0% in the original sample. The negative control region (NC; left/right ventricles) showed no significant supra-threshold voxels.

## 5 Discussion

We set out to improve the spatial specificity of detected regions using a “drill-down” approach—first selecting clusters, and then inferring on the voxels in those selected clusters—all with the same data. Reporting the proportion of active voxels in a cluster is an old quest in neuroimaging [e.g. Turkheimer et al., 2001], which also fulfills recent recommendations to report effect sizes, and not only active/inactive areas or cluster *p*-values [APA, 2001, Wilkinson, 1999].

Concerned with error inflation after the drill-down, we define an error to be an *over-estimation of the proportion of truly active voxels in a region*. Put differently, we are able to estimate the proportion of truly active voxels (PTD) in a selected cluster, with FWER control over clusters, all from the same data. This is made possible using results from Goeman and Solari [2011]. The fundamental observation is that if a statistical parametric map (SPM) of the brain satisfies the Simes inequality, then inferring on all possible voxel subsets is not hopelessly under-powered. Clearly, voxels within selected clusters are a subset of all voxel subsets, so that inference remains valid, for all possible subsets: in particular, for all the ones the practitioner queries after seeing the data.

Readers familiar with Scheffé’s post hoc test [Scheffe, 1953], may recognize that we use the same type of statistical reasoning. By controlling the error rate for all possible contrasts, Scheffé’s test allows the practitioner to choose the contrast after seeing the data. ARI does the same, not for all possible contrasts, but for all possible set selections. Moreover, and unlike Scheffé, ARI uses closed testing, which is more powerful in this context.

Quantifying the amount of true signal within clusters allows us to address the “spatial specificity paradox”, whereby the larger a cluster the less we know about the location of the signal. If the proportion of truly active voxels in a cluster is large, there is no real paradox. If this proportion is small, the practitioner should consider reducing the size of the clusters. Our two datasets demonstrate this is not a mere philosophical discussion, but rather, an empirical question with very real implications. The Go/No-go dataset shows small proportions of true activation within clusters, so that it is hard to tell which part of the cluster is truly active. The Auditory data shows large proportions, implying that the clusters are indeed mostly active.

The validation analysis shows consistent results across the two datasets: regions with a high PTD tend to have a high percentage of supra-threshold voxels in the validation dataset, and vice-versa. The PTD bounds are informative, i.e. not overly conservative, provided that the region is large enough: the gap between the PTD bound and the validated activation proportion is not too large. Smaller regions suffer a larger multiple testing burden, and so are more difficult to detect using ARI. The power of ARI is better for large clusters than for smaller ones, and better for small clusters than for single voxels. As a result it is easier to detect the presence of active voxels than it is to pinpoint them. A region may have a large proportion of active voxels, but we cannot usually say which voxels these are. This is especially true if the signal in a region is dispersed. In that case drilling down may not be successful: there may not always be a subregion with evidence for a larger PTD.

The ARI framework allows the practitioner a great deal of flexibility in that they can infer on regions, then drill down to voxels within regions, then redefine the regions, drill down in the new regions, etc. Users may iterate the process of choosing regions, bounding the PTD, and refining regions ad libitum, and without compromising FWER control. Since FWER control holds over all possible regions simultaneously, in fact any method for finding regions, using the same data or using external data, is allowed. Regions may be contiguous clusters, or any arbitrary, possibly disconnected, set of voxels. In particular, the ARI confidence bounds themselves may be used to select regions, and it is perfectly valid to select, for example, the largest region for which one is confident of a PTD of at least 0.7. While exploring the brain, computation time is not an issue. The underlying computations have been implemented in the R package hommel [Goeman et al., 2017], and take seconds to perform from *p*-value maps.

Our proofs assume the brain’s SPM satisfies the Simes inequality, while many analysis suites use a random field assumption (RFT) [Taylor and Worsley, 2007b]. The criticism of the validity of cluster inference voiced by Eklund et al. [2016] targets the RFT. We adopt the Simes inequality assumption because it facilitates our proofs, but it also means that if the ARI framework is used for cluster selection, it will not be subject to that criticism. The Simes inequality which we use instead of the RFT is implied by the PRDS condition, which has been established for brain SPMs by Nichols and Hayasaka [2003]. It may be possible to use RFT in combination with closed testing to obtain alternative lower bounds for PTD, but this is beyond the scope of this paper.

### 5.1 ARI for Cluster Selection

Our main innovation is in replacing the problem of cluster selection, with the problme of PTD-estimation. Cluster selection (i.e. testing) only claims that PTD > 0 for its regions, while ARI claims a non-trivial lower bound to PTD. Can ARI be used for the cluster selection itself? The answer is affirmative. The practitioner may toggle the cluster-wise cluster-forming threshold until reaching the desired PTD. E.g., select clusters with more than 70% of activation. Alternatively, the user may ask which clusters are the most active, i.e. where the activation is concentrated. Intuitively, the practitioner may “grow clusters”, i.e. decrease the cluster-forming threshold. At some point, the PTD will start to sharply decrease, and the practitioner would then stop growing the clusters. The *concentration set* in Meijer et al. [2016] formalizes this process. It gives a data-dependent *p*-value threshold above which ARI detects no signal. Used as a cluster-forming threshold, this gives a useful starting point from which to start the drill-down.

When using ARI for cluster selection, the question of power immediately arises. Naturally, the power properties of ARI are different to those of RFT-based models because the assumptions underlying the methods are different. RFT-selected clusters are not guaranteed to have *PTD* > 0 in ARI. However, such a comparison is not completely fair, as RFT-based methods only infer *PTD* > 0 on a limited number of clusters, while ARI infers the actual value of PTD on exponentially many clusters. Put differently, if a researcher only wishes to demonstrate *PTD* > 0 at a pre-chosen threshold, RFT based methods may be more powerful. If the researcher varies the cluster-forming threshold, focuses on sub-clusters, or wants to quantify the PTD, this is only possible with ARI.

Comparing power to other multiple testing procedures, we can say that ARI is strictly more powerful than voxel-wise FWER control. It guarantees *PTD* > 0 for every region containing a Bonferroni-significant voxel. For large regions it tends to give much larger values of PTD than would result from voxel-wise FWER.

A comparison of ARI to FDR-based methods is more complex because FDR is a much more relaxed criterion than FWER. Regarding voxel-wise FDR control, it holds that whenever the Benjamini-Hochberg algorithm [Benjamini and Hochberg, 1995] detects at least one active voxel, ARI will find at least one region with *PTD* > 0. Direct comparison with cluster-wise FDR is impossible, especially since there is at this moment no method that controls FDR over all clusters as ARI does.

## 6 Methods

### 6.1 Preprocessing

Acquisition parameters and detailed information about the stimuli of both datasets can be found in Lee et al. [under review] and Pernet et al. [2015]. Both datasets were analyzed in FSL [Jenkinson et al., 2012] using a standard preprocessing pipeline. Time-series data was high-pass filtered (Go/No-go, 90 seconds; Auditory, 128 seconds). Functional images were brain extracted [Smith, 2002], spatially smoothed (6 mm full width at half maximum), and registered to standard space using linear registration (FLIRT [Jenkinson and Smith, 2001, Jenkinson et al., 2002] with 12 degree-of-freedom boundary-based registration). Six motion regressors (MCFLIRT [Jenkinson et al., 2002]) and periods with excessive motion were modeled as additional confound regressors. Boxcar functions of the stimulus timings for the different conditions were convolved with a double-gamma hemodynamic response function, with a temporal derivative to model differences in slice acquisition time. For the Go/No-go dataset we analyzed the No-go > Go contrast using FEAT [Woolrich et al., 2001], using FLAME 1 estimation with a cluster-threshold multiple comparison correction based on RFT, highlighting brain regions involved in successfully inhibiting a response. For the auditory dataset we analyzed the Vocal > Non-vocal contrast, highlighting brain regions involved in speech processing.

ARI was performed on both our datasets using our own implementation, made publicly available in the hommel package [Goeman et al., 2017] for the R software environment [Team, 2000]. We used the analysis pipeline described in Section 3.5.

### 6.2 ARI Validation

If the PTD of a cluster is bounded by *q*, then in a new dataset, a proportion of at least *q* of the voxels in the cluster are true signal, and thus should be rediscovered. To validate that this is indeed the case, we used the clusters from the Vocal > Non-vocal contrast of the first set of 33 subjects, and computed the proportion of supra-threshold voxels (uncorrected *p*-value < .05) in a new set of 66 subjects.

Table 2 for the Auditory dataset therefore includes additional columns for the number and percentage of suprathreshold voxels in the verification dataset. In addition, we calculated the PTD in a negative control region where we don’t expect any significant voxels. We used the left and right ventricles (Harvard-Oxford atlas with a probability threshold > 50%), and calculated the PTD in each of these areas for both Auditory datasets.

## Footnotes

1 Sometimes known as *multi-resolution*, or *post hoc*.

2 Readers familiar with the *false detection rate* literature, will note that *PTD* = 1 — *FDP*, where *FDP* is the *false discovery proportion*.

